# Phenanthrene contamination and ploidy level influence the rhizosphere microbiome of *Spartina*

**DOI:** 10.1101/625657

**Authors:** Armand Cavé-Radet, Cécile Monard, Abdelhak El-Amrani, Armel Salmon, Malika Ainouche, Étienne Yergeau

## Abstract

*Spartina* spp. are widely distributed salt marsh plants that have a recent history of hybridization and polyploidization. These evolutionary events have resulted in species with a heightened resilience to hydrocarbon contamination, which could make them an ideal model plant for the phytoremediation/reclamation of contaminated coastal ecosystems. However, it is still unknown if allopolyploidization events also resulted in differences in the plant rhizosphere-associated microbial communities, and if this could improve the plant phytoremediation potential. Here, we grew two parental *Spartina* species, their hybrid and the resulting allopolyploid in salt marsh sediments that were contaminated or not with phenanthrene, a model tricyclic PAH. The DNA from the rhizosphere soil was extracted and the bacterial 16S rRNA gene and ITS region were amplified and sequenced. Generally, both the presence of phenanthrene and the identity of the plant species had significant influences on the bacterial and fungal community structure, composition and diversity. In particular, the allopolyploid *S. anglica*, harbored a more diverse bacterial community in its rhizosphere, and relatively higher abundance of various bacterial and fungal taxa. Putative hydrocarbon degraders were significantly more abundant in the rhizosphere soil contaminated with phenanthrene, with the *Nocardia* genus being significantly more abundant in the rhizosphere of *S. anglica*. Overall our results are showing that the recent polyploidization events in the *Spartina* did influence the rhizosphere microbiome, both under normal and contaminated conditions, but more work will be necessary to confirm if these differences result in a higher phytoremediation potential.

**Importance:** Salt marshes are at the forefront of coastal contamination events caused by marine oil spills. Microbes in these environments play a key role in the natural attenuation of these contamination events, often in association with plant roots. One such plant is the Spartina, which are widely distributed salt marsh plants. Intriguingly, some species of the Spartina show heightened resistance to contamination, which we hypothesized to be due to differences in their microbiota. This was indeed the case, with the most resistant Spartina also showing the most different microbiota. A better understanding of the relationships between the Spartina and their microbiota could improve the coastal oil spill clean-up strategies and provide green alternatives to more traditional physico-chemical approaches.

## Introduction

Polycyclic aromatic hydrocarbons (PAHs) are a group of ubiquitous organic pollutants, that are highly concerning in view of their potentially severe impact on natural ecosystems and public health. The remediation of sites contaminated by these compounds due to human activity, such as oil spill, is mainly carried out through excavation, soil leaching or various techniques based on microbial degradation (1, 2). Phytoremediation is an alternative low-cost and environmentally-friendly technology that uses plants and their associated microorganisms to remove pollutants from the environment. Phytoremediation exists under various flavors depending on the type of contaminant targeted and its fate (3). For instance, phytoextraction mainly concerns the extraction of inorganic pollutant from the soil and its translocation and accumulation in plant tissues. Volatile compounds can also be extracted from the soil by plants, and after translocation be released into the atmosphere during phytovolatilization. However, one of the most interesting form of phytoremediation is rhizodegradation, where plants stimulate a wide variety of root-associated bacteria and fungi (4) to degrade organic contaminants, often all the way to CO_2_ (mineralization). Clearly, rhizodegradation is a plant-microbe joint venture. Close interactions between plants and soil microorganisms were reported to play a major role in detoxification and metabolization of xenobiotics (5, 6), and recent advances in ‘omics’ approaches now allow to address the plant-microbe complex in its entirety (7). Root exudates were demonstrated to have a central role, modifying the microbial PAH degraders community structure (8, 9) and significantly increasing microbial biomass (10). Rhizodegradation of PAH has even been suggested to be a model for understanding plant-microbe interactions (11).

One area of interest that is understudied is the use of phytoremediation for the cleanup of coastal salt marshes contaminated by hydrocarbons coming from seawater spills. In that context, *Spartina* is a particularly interesting genus, colonizing salt marshes all around the world and providing key ecosystem services, while being particularly resilient to the effects of oil spills (12–15). This genus is also characterized by numerous hybridization and genome doubling events (polyploidy) (16), which are major evolutionary mechanisms for eukaryotes, and most specially for plants (17, 18). Recent work from our group has shown that this genome doubling increased tolerance to the model PAH phenanthrene in *S. anglica* as compared to its single-genome parents (19). Indeed, the genomic shock induced by the merging of divergent genomes results in massive shifts in genetic and epigenetic pathways, that may lead to the emergence of new adaptive phenotypes to environmental constraints (20). Although previous studies have shown a link between plant genotype and the rhizosphere microbial community structure and gene expression during phytoremediation (21, 22), nothing is known about the effects of genome doubling events on the roots associated microorganisms and their response to PAH contamination. Here, we hypothesized that the heightened resistance of polyploid *Spartina* species to PAH contaminants is partly related to concomitant shifts in the root-associated microorganisms. We selected four *Spartina* species: the parental species *Spartina alterniflora* (2n=6x=62) and *S. maritima* (2n=6x=60), their interspecific sterile hybrid *S. x townsendii* (2n=6x=62) that appeared and emerged at the end of the 19^th^ century and the allopolyploid derivative *S. anglica* that subsequently appeared following genome doubling (2n=12x=120, 122, 124). These different *Spartina* species were grown in salt marsh sediments contaminated or not with phenanthrene, after which the bacterial 16S rRNA gene and fungal ITS region of the rhizosphere soil were amplified and sequenced.

## Material and methods

### Plant and soil material

Plants were collected in 2016 on their natural habitats along coastlines of France and England. For the parental species, *S. alterniflora* was sampled at Le Faou (Roadstead of Brest, France), whereas *S. maritima* was sampled in Brillac-Sarzeau and Le Hezo (Gulf of Morbihan, France). The homoploid hybrid *S. x townsendii*, which distribution range is not extended to France was collected in Hythe (Romney Marsh, England), and the allopolyploid *S. anglica* was sampled at La Guimorais (Saint-Coulomb, France), where bulk sediments (not associated with plants) were also collected. All the plants were acclimated in the La Guimorais sediments for three weeks before the start of the experiment. For the experiment, the remaining sediments were rinsed several times, dried and sieved through a 5mm sieve. To contaminate the soil, 150 g of air-dried sediment were either spiked with 45 mg of phenanthrene diluted in 10 mL of absolute ethanol, whereas the controls were spiked with 10 mL of absolute ethanol. After total evaporation of ethanol for one day, these sediment samples were vigorously mixed by hand with an additional 150 g of air-dried sediment to reach a final concentration of 150 mg phenanthrene kg^−1^ substrate for the contaminated treatment. The contaminated and the control soils were sampled at this step and kept at −20°C until DNA extraction.

### Experimental design

Both polluted and control sediments were supplemented with sterile vermiculite before the start of the experiment (volume 1/3) for a better substrate breathability. Individual *Spartina* plants (four species) were then transplanted in pot containing 3 kg of treated or control substrates, and rhizomes were carefully placed into rhizobags of 3 cm of diameter, for a physical separation of the rhizosphere. The experimental design was replicated three times resulting in a total of 24 pots that were placed in a phytotronic chamber with a light/dark regime of 16/8h, in an average ambient temperature of 20°C. Pots were watered with 200 mL of half-strength Hoagland’s nutrient solution (23) every five days. After 60 days of growth, plants were uprooted and the rhizosphere soil (inside the rhizobags) was collected and stored at −20°C until DNA extraction.

### Bacterial 16S rRNA gene and fungal ITS amplification

Genomic DNA from soil samples was extracted using a MoBio PowerSoil DNA extraction kit. Amplicons were prepared from total DNA by PCR targeting the bacterial 16S rRNA (forward: 515F: GTGCCAGCMGCCGCGGTAA and reverse: 806R: GGACTACHVGGGTWTCTAAT) (24) and fungi ITS (forward: ITS1F: CTTGGTCATTTAGAGGAAGTAA and reverse: 58A2R: CTGCGTTCTTCATCGAT) (25) specific primers. PCR amplification were performed in 25 μL final reaction volumes containing final concentrations of 1X KAPA HiFi HotSart ReadyMix, 0.4 mg.ml^−1^ bovine serum albumin, 0.6 μM forward and reverse primers, and sterile water and 1 μL of DNA. PCR conditions were as followed: 5 min of initial denaturation (95°C), 25 cycles of 30s denaturation (95°C), 30s for primer annealing (55°C), and 45s elongation (72°C), followed by final extension for 10 min at 72°C. PCR amplicons were then purified using AMPure XP beads. Amplicon indexing was carried by a second PCR in 25 μL final reaction volumes containing 12.5 μL 2X KAPA HiFi HotSart ReadyMix, 2.5 μL of specific index primers 1 and 2 from the Nextera Index kit, and 5 μL of purified amplicon. The second step PCR conditions were as followed: 3 min of initial denaturation (95°C), 8 cycles of 30s denaturation (95°C), 30s for primer annealing (55°C), and 45s elongation (72°C), followed by final extension for 5 min at 72°C. Index PCR cleanup was performed using AMPure XP beads. Finally, PCR amplicons were quantified using PicoGreen, normalized at 1 ng μl^−1^, pooled together and sent for sequencing on an Illumina MiSeq (paired-end 2×250bp) at the McGill University and Genome Quebec Innovation Center (Montréal, Canada). Raw reads and associated metadata are available through NCBI BioProject accession PRJNA518897 (http://www.ncbi.nlm.nih.gov/bioproject/518897).

### Sequencing data processing

Raw sequencing reads were processed using the mothur MiSeq Standard Operating Procedure (SOP) (26). The bacterial 16S rRNA gene and fungal ITS reads were processed separately. Bacterial paired-end reads were first assembled and contigs with ambiguous bases or smaller than 275 bp were excluded. Fungal paired reads were assembled, but because ITS amplicons have variable lengths, only the contigs with ambiguous bases were filtered out (no exclusion based on contig size). The dataset was dereplicated for faster computation, after which chimera were removed. Unique bacterial sequences were aligned using the SILVA nr database (v128) after which the aligned 16 rRNA gene and ITS sequences were clustered at 97% sequence similarity, and singletons were removed from the analysis. Bacterial sequences were classified with the 16S rRNA PDS reference from Ribosomal Database Project (RDP version 16; 80% cut-off on bootstrap value for confidence taxonomic assignment) and undesirable taxonomic assigned sequences (chloroplast, mitochondria, unknown, Archaea and Eukaryota) were removed. Unique fungal sequences were classified with UNITE ITS database (99% singletons provided on https://mothur.org/wiki/UNITE_ITS_database; 80% cut-off) and undesirable taxonomic assigned sequences (unknown, Plantae and Protista) were removed. Misannotated fungal OTU (Operational Taxonomic Unit) corresponding to *Spartina* ITS sequences were identified using BLASTn (27) alignments on GenBank rDNA *Spartina* public data (best blast hit, min 90% identity and 80% query coverage) and removed from the analysis. Totals of 4,905,456 and 808,608 sequences were respectively assembled from bacterial 16S rRNA gene and fungal ITS reads. Processing of bacterial 16S rRNA gene data resulted in 3,132,223 sequences, corresponding to 121,804 unique sequences that clustered into 28,092 OTUs. For the ITS sequences, we retained 785,825 sequences corresponding to 538,892 unique sequences that clustered into 10,550 OTUs. Total of 38 misannotated OTUs corresponding to *Spartina* ITS sequences were removed from the analysis, resulting into 10,512 clean fungi OTUs.

### Statistical analyses

Data and statistical analyses were performed in R (v 3.5.1). Indices of α-diversity (inversed Simpson index) related to bacterial and fungal OTUs were calculated after normalization through random subsampling based on the size of the smallest library. Comparisons between α-diversity of microbial communities, the relative abundance of phyla and the relative abundance of selected putative hydrocarbon degraders according to the *Spartina* species and the treatments (polluted or control substrate) were conducted using ANOVA with post-hoc Tukey HSD tests (aov and tukeyHSD functions of the *stats* package). Principal coordinate analysis (PCoA) to describe β-diversity was performed on normalized OTU tables using Bray-Curtis dissimilarity index in the vegan (28) and ape (29) packages. Permanova testing the impact of *Spartina* species, treatments, and their interactions were conducted through 999 permutations. Indicative species among OTUs were investigated using default parameters provided by the indicspecies package (30).

## Results

### Bacterial and fungal diversity

The presence of phenanthrene led to a general decrease of bacterial diversity in the rhizosphere of all the plant species (two-way ANOVA: F = 12.41, *P* = 0.002; Fig. 1a) but was only significant for *S. anglica* (pairwise tests; Fig. 1a). The two-way ANOVA also highlighted a significant effect of plant species on the bacterial diversity (F = 6.19 *P* = 0.005), and Tukey HSD post-hoc tests showed that the rhizosphere of *S. anglica* was significantly more diverse than the rhizosphere of *S. alterniflora* and *S. maritima*. Pairwise tests confirmed this trend, with the bacterial diversity in the control rhizosphere of *S. anglica* being higher than in the control rhizosphere of *S. alterniflora* and in all contaminated rhizospheres (Fig. 1a). The interaction term Phenanthrene x Species did not have a significant effect on the bacterial diversity in two-way ANOVA tests. For fungi, phenanthrene contamination and the interaction term did not have a significant effect on the diversity in the rhizosphere in two-way ANOVA tests. However, plant species did have a significant effect on the fungal diversity in the rhizosphere (F = 11.20, *P* < 0.001), and Tukey HSD post-hoc test showed that *S. alterniflora* harbored a significantly more diverse fungal community in its rhizosphere as compared to the other three species (Fig. 1b).

**Figure 1.**
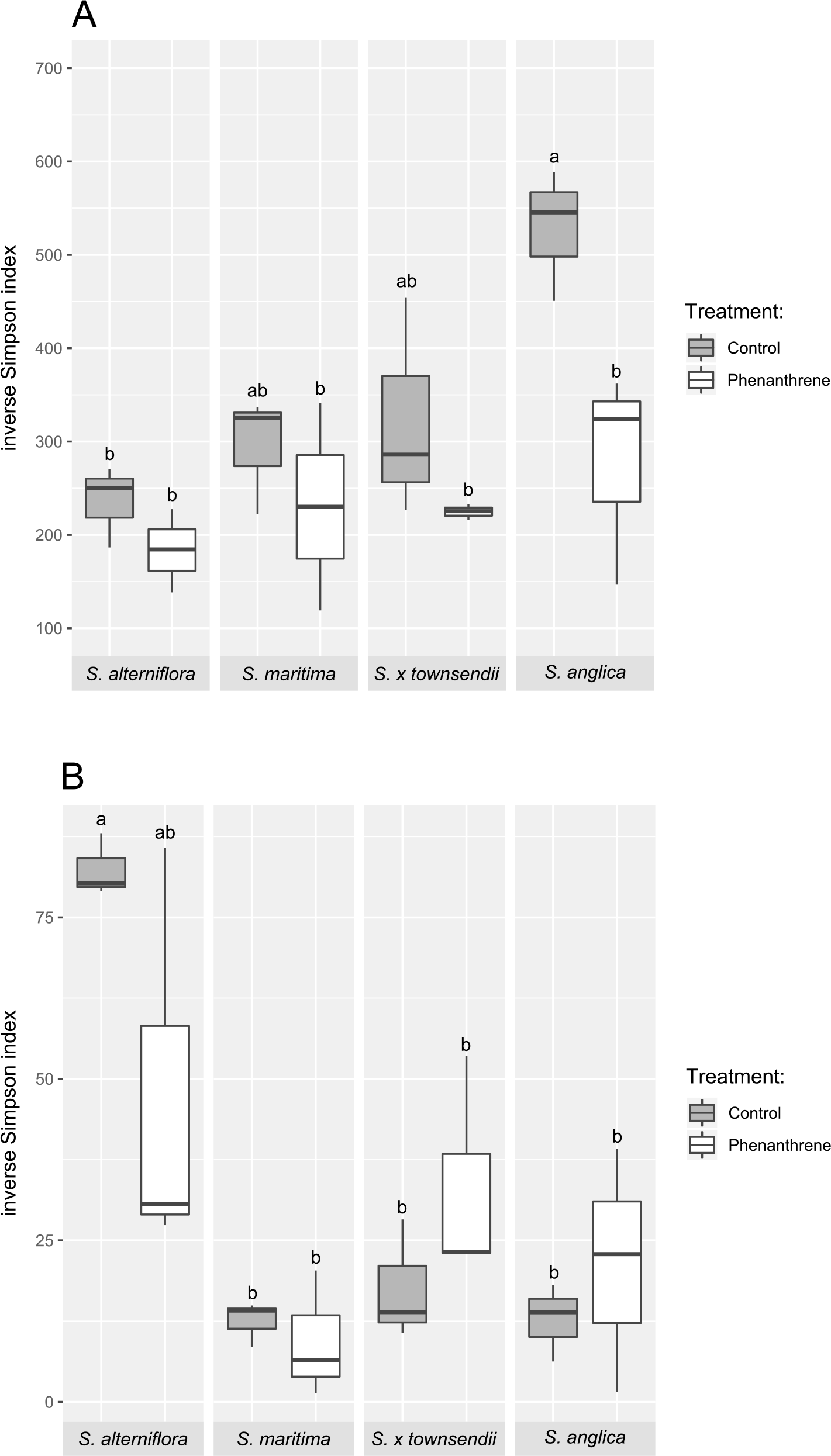
(A) Bacterial and (B) fungal diversity (inverse Simpson index) in the rhizosphere of four different *Spartina* species grown in phenanthrene-contaminated or control sediments. Values annotated by different letters are significantly different according to pairwise t-ests (Welch correction and Bonferroni adjusted, *P* < 0.05).

### Bacterial and fungal community structure

Principal coordinate analysis (PCoA) ordinations based on bacterial OTUs showed some level of clustering based on plant species and contamination (Fig. 2a). Permanova tests confirmed that *Spartina* species (F = 2.09, *P* < 0.001) and phenanthrene (F = 2.48, *P* < 0.001) had highly significant influences on the bacterial community structure, with a slightly stronger effect for contamination (higher F-ratio). For fungi, the PCoA ordination showed a relatively clearer picture than bacteria, with clustering of many treatments and a separation of most samples based on the contamination treatment along the second axis (Fig. 2b). Permanova confirmed the slightly higher explanatory power of the contaminant treatment (F = 1.94, *P* < 0.001) as compared to the plant species (F = 1.67, *P* < 0.001). Nevertheless, both factors were highly significant, and even the interaction term was significant (F = 1.32, *P* = 0.018), suggesting that the contamination did not affect the rhizosphere fungal community structure in the same way for all plant species.

**Figure 2.**
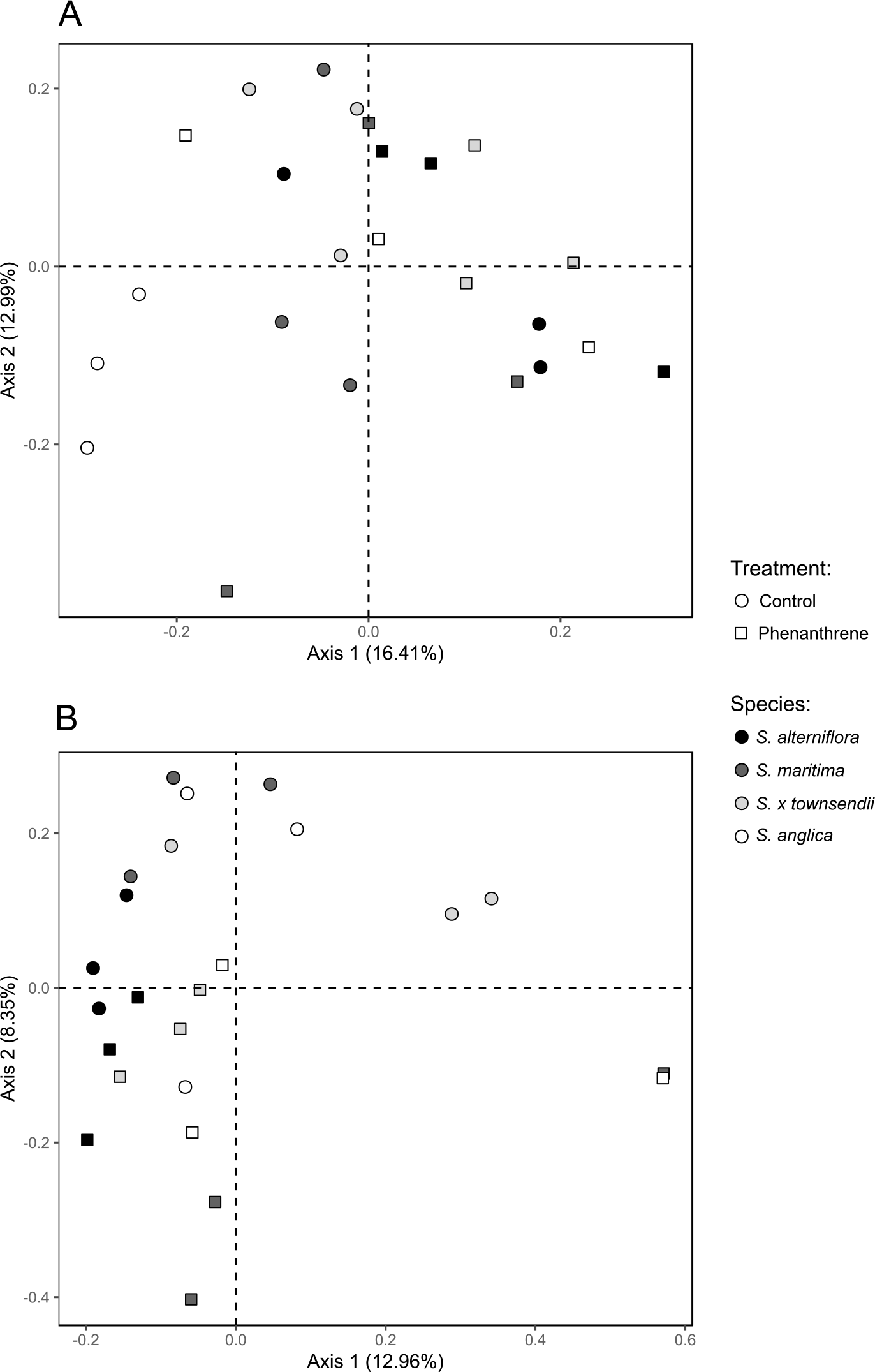
Principal coordinate analysis (PCoA) for the (A) bacterial and (B) fungal communities of the rhizosphere of four different *Spartina* species grown in phenanthrene-contaminated or control sediments.

### Bacterial and fungal community composition

The relative abundance of various dominant bacterial phyla in the rhizosphere of *Spartina* varied between the different plant species and between the contaminated and non-contaminated soils (Fig. 3). These shifts were not significant for the *Acidobacteria*, the *Actinobacteria*, the *Bacteroidetes*, and the *Alpha*-, *Beta*-, *Delta*- and *Epsilonproteobacteria*. In contrast, plant species significantly influenced the rhizosphere relative abundance of *Chloroflexi* (F = 6.21, *P* = 0.05, significantly higher in the rhizosphere of *S. anglica* as compared to *S. alterniflora* and *S. x townsendii* in Tukey HSD tests), *Planctomycetes* (F=8.86, P=0.00108, significantly higher in the rhizosphere of *S. anglica* as compared to *S. alterniflora* and *S. x townsendii,* and in the rhizosphere of *S. maritima* vs. *S. alterniflora* in Tukey HSD tests) and total *Proteobacteria* (F = 4.54, *P* = 0.02, significantly lower in the rhizosphere of *S. anglica* as compared to *S. alterniflora* and *S. x townsendii* in Tukey HSD tests). A significant effect of contamination was also observed for the *Chloroflexi* (F = 20.93, *P* < 0.001), total *Proteobacteria* (F = 14.23, *P* = 0.002), *Verrucomicrobia* (F = 4.85, *P* = 0.43) and *Gammaproteobacteria* (F = 11.39, *P* = 0.004). The genera *Sphingobacterium*, *Acinetobacter*, *Nocardia*, *Pseudomonas*, *Mycobacterium*, *Burkholderia*, *Bacillus*, *Sphingomonas*, *Rhodococcus*, *Paenibacillus*, *Massilia, Alcanivorax, Cycloclasticus* were singled out as putative hydrocarbon degraders based on a survey of the available literature. The summed relative abundance of all these genera varied between around 2% to over 3% of all reads and was significantly higher in the rhizosphere of plants growing in contaminated soil (F = 8.43, *P* = 0.01, Fig. 4), but showed no significant differences between plant species. For the individual genera, the relative abundance of *Paenibacillus, Bacillus, Burkholderia, Acinetobacter, Alcanivorax, Rhodococcus* and *Pseudomonas* did not vary significantly between the plant species and treatments. However, the relative abundances of *Sphingomonas* (F = 9.85, *P* < 0.001, significantly higher in the rhizosphere of *S. x towsendii* as compared to all other species in Tukey HSD tests), *Sphingobacterium* (F = 5.41, *P* = 0.009, significantly higher in the rhizosphere of *S. maritima* as compared to all other species in Tukey HSD tests), *Nocardia* (F = 7.56, *P* = 0.002; significantly higher in the rhizosphere of *S. anglica* as compared to all other species in Tukey HSD tests) showed significant differences between the rhizosphere of the different plant species. The relative abundances of *Massilia* (F = 17.54, *P* < 0.001), *Cycloclasticus* (F = 4.96, *P* = 0.04) and *Mycobacterium* (F = 7.49, *P* = 0.01) were significantly affected by the phenanthrene treatment, with significantly higher relative abundance in the rhizosphere of plants growing in the contaminated soil (Fig. 4). In addition, the interaction term was significant for *Sphingobacterium* (F = 4.73, *P* = 0.001). Indicator species analyses confirmed some of the trends observed in ANOVA tests, as it identified bacterial OTUs related to *Massilia* and *Cycloclasticus* as the OTUs with the highest indicator power for contaminated rhizospheres (Table 1).

**Figure 3.**
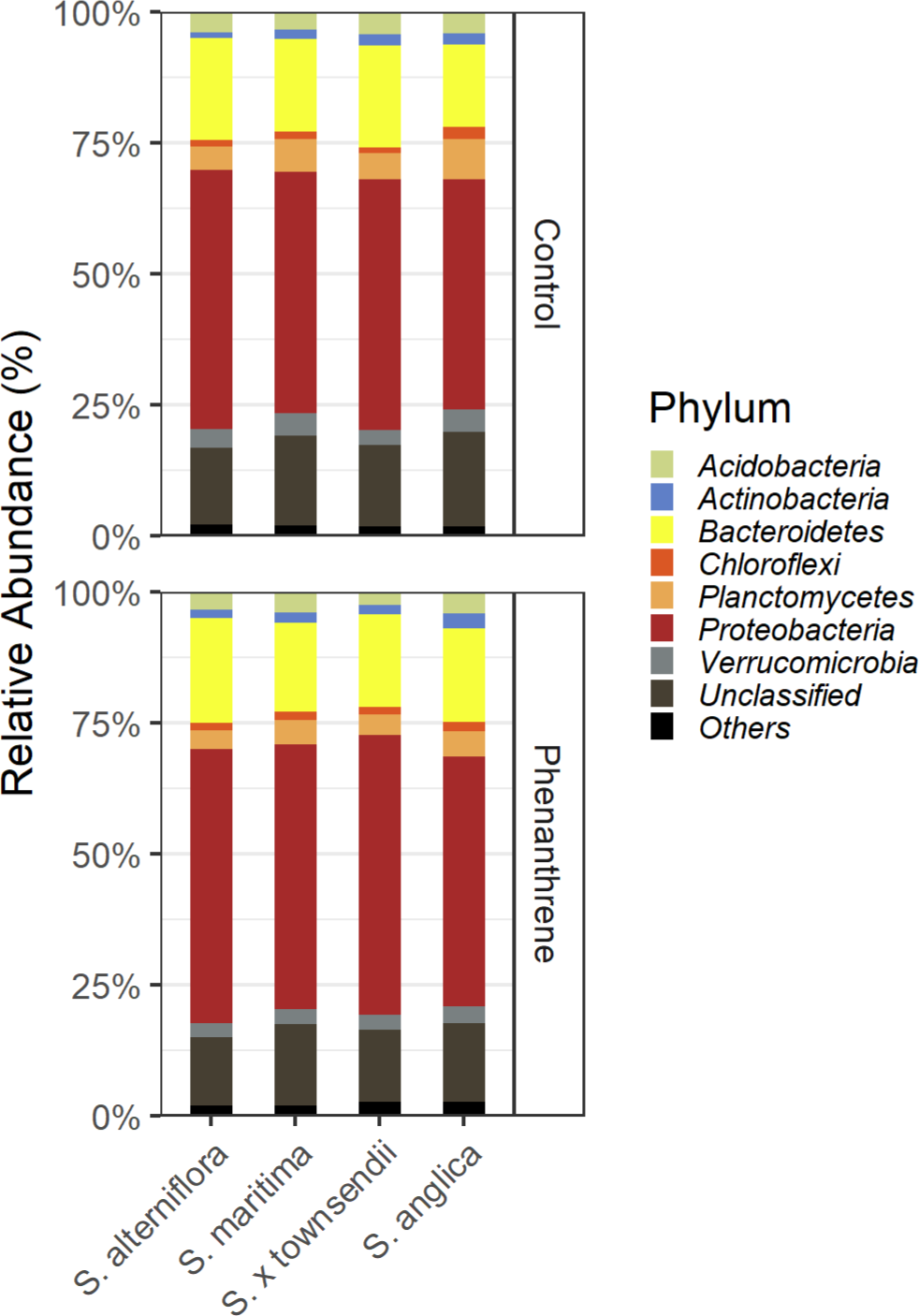
Relative abundance of the most represented bacterial phyla in the rhizosphere of four different *Spartina* species grown in phenanthrene-contaminated or control sediments.

**Figure 4.**
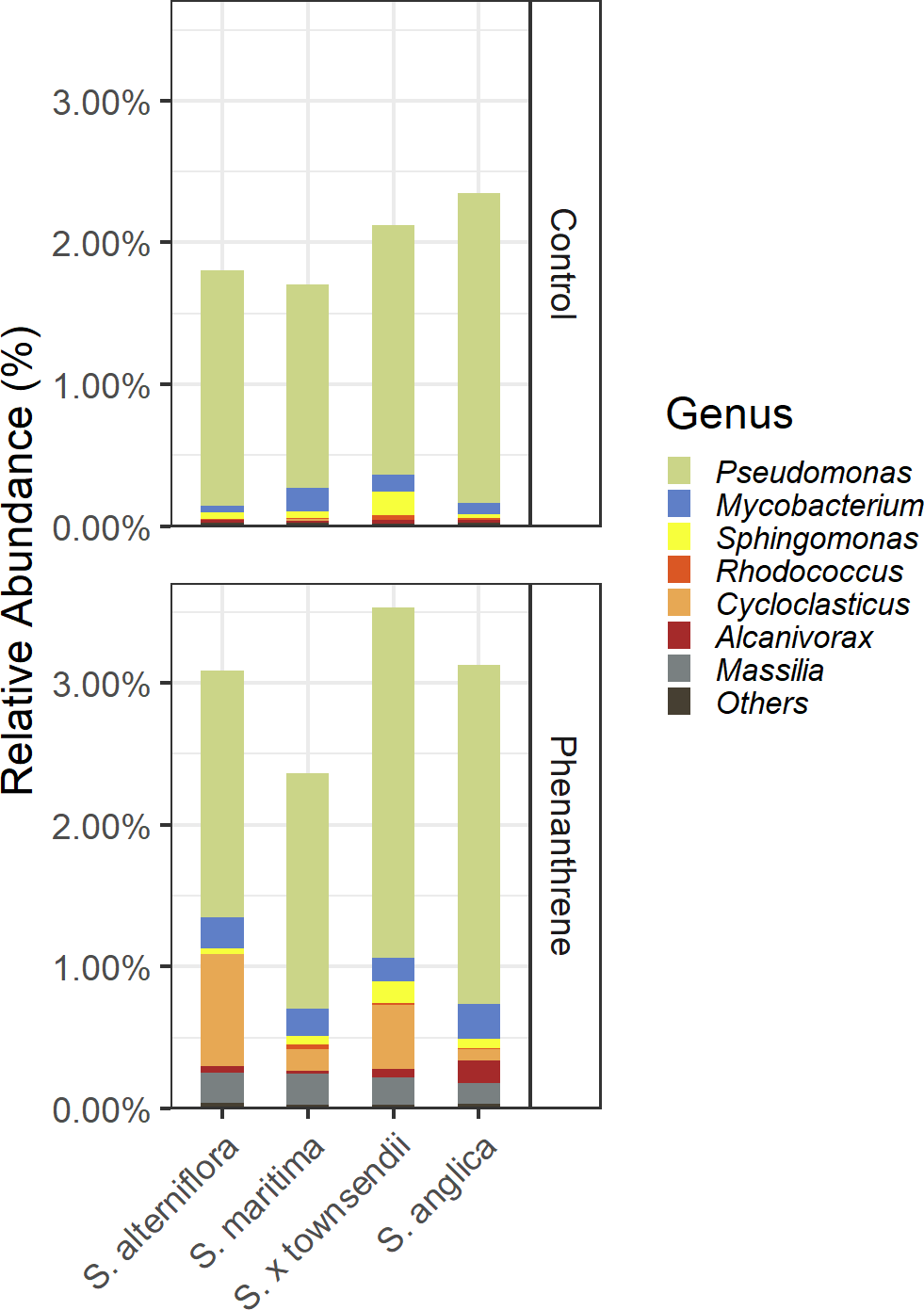
Relative abundance of putative PAH-degrading bacteria genera in the rhizosphere of four different *Spartina* species grown in phenanthrene-contaminated or control sediments.

**Table 1.** Indicator bacterial OTUs for the phenanthrene-contaminated *Spartina* rhizospheres.

| OTU | stat | p.value | Assigned genera |
| --- | --- | --- | --- |
| Otu00188 | 0.998 | 0.001 | *Massilia* |
| Otu00140 | 0.996 | 0.001 | *Cycloclasticus* |
| Otu00370 | 0.991 | 0.001 | *Alcaligenaceae* unclassified |
| Otu00552 | 0.989 | 0.049 | *Flavobacteriaceae* unclassified |
| Otu02955 | 0.960 | 0.002 | *Salinirepens* |
| Otu01642 | 0.954 | 0.006 | *Sphingobium* |
| Otu02528 | 0.954 | 0.005 | *Alcaligenaceae* unclassified |
| Otu01089 | 0.941 | 0.005 | *Peredibacter* |
| Otu04988 | 0.921 | 0.008 | unassigned |
| Otu03519 | 0.896 | 0.006 | *Gammaproteobacteria* unclassified |
| Otu08173 | 0.859 | 0.028 | unassigned |
| Otu02835 | 0.853 | 0.026 | unassigned |
| Otu04910 | 0.841 | 0.037 | *Bacteroidetes* unclassified |

For fungi, the relative abundance of the three main phyla detected in the rhizosphere, varied between the plant species and the treatments (not shown). However, these trends were only significant for the *Basidiomycota*, where the plant species (F = 4.14, *P* = 0.02), the contamination treatment (F = 6.17, *P* = 0.02) and the interaction term (F = 4.62, P = 0.02) were all significant. However, no fungal OTUs were identified as indicator of phenanthrene contamination.

## Discussion

The hypothesis behind the present study was that the increased resilience of the allopolyploid *Spartina anglica* to hydrocarbon contamination was partly due to its root-associated microbial communities. The results partly confirmed this trend, as we found significant differences between the different *Spartina* species in terms of their bacterial and fungal community composition, structure and diversity. The allopolyploid *S. anglica* harbored a more diverse bacterial community, composed of relatively more *Nocardia*, *Chloroflexi* and *Planctomycetes* and less *Proteobacteria* in its rhizosphere as compared to its diploid parents and their hybrid. Previous studies from our group using willows have shown that the phylogeny of the host plant significantly influenced the fungal community composition, but only under high levels of contaminant (21). We also showed that the metatranscriptomic response of the rhizosphere microbial communities to contamination varied between different willow genotypes, and that this response was mirrored in the growth of the genotypes in contaminated soil (22). However, this is the first time, to our knowledge, that differences between the rhizosphere microbiome of recently naturally speciated plants are reported in the context of soil contamination. One particularly interesting aspect of these differences is the higher bacterial diversity in the rhizosphere of *S. anglica* as compared to the other plant species. Upon a contamination event, the plant-associated microbial diversity could have a central importance, as higher diversity generally results in functional redundancy, which would enable the microbiome to cope with a wider variety of environmental conditions while still providing essential services to the plant. Previous studies from our group have shown that the initial soil diversity explained better the difference in willow growth under highly contaminated conditions than diversity at the end of the experiment (31). It was further suggested that restoring microbial diversity of degraded environments could be the key for successful phytoremediation. Even though we cannot exclude the possibility that the higher diversity in the rhizosphere of *S. anglica* is because the bulk soil used for the experiment was from a site where *S. anglica* predominantly grew, this link between the higher rhizosphere diversity and the increased resilience of *S. anglica* to contamination is intriguing and warrants further research.

The rhizosphere microbial communities of the four *Spartina* species tested showed many common responses to the presence of the contaminant. In the presence of PAH, salt marsh plants were reported to harbor different microbial communities, favoring PAH degrading microorganisms (32). Similarly, we found here that the general decrease in bacterial diversity under phenanthrene was concomitant to an increase in the relative abundance of putative PAH degraders, and more specifically for the genera *Mycobacterium*, *Cycloclasticus* and *Massilia*. These genera represent large bacterial groups that are able to metabolize and degrade PAH (33, 34) and are consistent with previous results about the PAH-degrading bacteria associated to *Spartina* in salt marshes (35, 36). Interestingly, *Cycloclasticus* spp. were often reported as one of the predominant PAH degraders in seawater (37, 38), whereas *Mycobacterium* are typical soil PAH degraders (39) and *Massilia* were found to degrade PAHs in soils or associated to roots (40, 41). This diversity of putative PAH degraders associated with the rhizosphere of *Spartina* is probably linked to the nature of its habitat, at the interface of terrestrial, plant and maritime ecosystems. These bacterial genera probably contain candidates of interest for salt marsh remediation, but their potential would have to be confirmed by other complementary methods.

We described here for the first time the root-associated microbiome of four *Spartina* species in a context of allopolyploidization and PAH contamination. Significant differences were observed between the plant species and between contaminated and control rhizospheres. Admittingly, shifts in the relative abundance and diversity of bacteria and fungi taxa might not be the only factor contributing to the increased resilience of *S. anglica* to contaminant stress, and the shifts in the expression of specific genes or changes in the community size could be important and not captured by amplicon sequencing. More work would be required to confirm if the differences observed here are linked to the increased resilience of the allopolyploid species in the face of hydrocarbon contamination.

## Acknowledgements

A.C.-R. was supported by an Erasmus+ mobility scholarship from the European Commission. This work was supported by the Ministère de l’Enseignement Supérieur et de la Recherche, by the CNRS, the Observatoire des Sciences et de l’Univers de Rennes (OSUR) and by a NSERC Discovery Grant (2014-05274 to EY). This research was enabled in part by support provided by Calcul Québec (www.calculquebec.ca) and Compute Canada (www.computecanada.ca).

